# Implications of central carbon metabolism in SARS-CoV-2 replication and disease severity

**DOI:** 10.1101/2021.02.24.432759

**Authors:** Shuba Krishnan, Hampus Nordqvist, Anoop T. Ambikan, Soham Gupta, Maike Sperk, Sara Svensson-Akusjärvi, Flora Mikaeloff, Rui Benfeitas, Elisa Saccon, Sivasankaran Munusamy Ponnan, Jimmy Esneider Rodriguez, Negin Nikouyan, Amani Odeh, Gustaf Ahlén, Muhammad Asghar, Matti Sällberg, Jan Vesterbacka, Piotr Nowak, Ákos Végvári, Anders Sönnerborg, Carl Johan Treutiger, Ujjwal Neogi

**Affiliations:** Division of Clinical Microbiology, Department of Laboratory Medicine, Karolinska Institute, ANA Futura, Campus Flemingsberg, S-14152 Stockholm, Sweden; Södersjukhuset (The South General Hospital), Stockholm, Sweden; National Bioinformatics Infrastructure Sweden (NBIS), Science for Life Laboratory, Department of Biochemistry and Biophysics, Stockholm University, S-10691 Stockholm, Sweden; Centre for Infectious Disease Research, Indian Institute of Science (IISc), CV Raman Avenue, Bangalore, India; Division of Chemistry I, Department of Medical Biochemistry and Biophysics, Karolinska Institutet, SE-171 65 Stockholm, Sweden; Division of Infectious Diseases, Department of Medicine Solna, Karolinska Institutet, Stockholm, Sweden; Department of Medicine Huddinge, Division of Infectious Diseases, Karolinska Institute, I73, Karolinska University Hospital, Huddinge, 141 86 Stockholm, Sweden; The Laboratory for Molecular Infection Medicine Sweden MIMS, Umeå University, Sweden.

## Abstract

Viruses hijack host metabolic pathways for their replicative advantage. Several observational trans-omics analyses associated carbon and amino acid metabolism in coronavirus disease 2019 (COVID-19) severity in patients but lacked mechanistic insights. In this study, using patient-derived multi-omics data and *in vitro* infection assays, we aimed to understand i) role of key metabolic pathways in severe acute respiratory syndrome coronavirus-2 (SARS-CoV-2) reproduction and ii) its association with disease severity. Our data suggests that monocytes are key to the altered immune response during COVID-19. COVID-19 infection was associated with increased plasma glutamate levels, while glucose and mannose levels were determinants of the disease severity. Monocytes showed altered expression pattern of carbohydrate and amino acid transporters, GLUT1 and xCT respectively in severe COVID-19. Furthermore, lung epithelial cells (Calu-3) showed a strong acute metabolic adaptation following infection *in vitro* by modulating central carbon metabolism. We found that glycolysis and glutaminolysis are essential for virus replication and blocking these metabolic pathways caused significant reduction in virus production. Taken together, our study highlights that the virus utilizes and re-wires pathways governing central carbon metabolism leading to metabolic toxicity. Thus, the host metabolic perturbation could be an attractive strategy to limit the viral replication and disease severity.

## Introduction

The global pandemic of coronavirus disease 2019 (COVID-19) caused by the severe acute respiratory syndrome coronavirus-2 (SARS-CoV-2) created a severe public health crisis worldwide. Although most patients presented with mild to moderate or no symptoms, patients having pre-existing metabolic disorders like diabetes, cardiovascular diseases, and obesity are at risk for severe and critical cases of infection. Some recent observational studies indicate that disease severity in COVID-19 patients are associated with plasma metabolic abnormalities that include a shift towards amino acid and fatty acid synthesis, altered energy and lipid metabolism [1–3] and metabolic profile can identify the disease severity [4]. However, metabolic regulation of an individual always depends on several factors including age, gender, environmental factors, dietary intake and lifestyle. Such alterations in metabolic regulation can change rapidly or adapt to an altered situation and sometimes have sustained effects over an extended period. The initial phase of characterization of the metabolic landscape of COVID-19 and its association with disease severity has urged the need to understand how metabolic reprogramming occurs during the acute SARS-CoV-2 infection with the ultimate goal towards therapeutic intervention.

Viruses are known to exploit the host metabolic machinery to meet their biosynthetic demands for optimal replication capacity. This cellular exploration is highly connected with the initial host-viral response, thereby determining the disease pathogenesis. Viral replication is dependent on extracellular carbon sources such as glucose and glutamine. It induces a plethora of metabolic alterations in host-cell including host central carbon metabolism, nucleotide, fatty acids and lipid synthesis that modulate viral pathogenesis and host-response [5, 6]. Recent *in vitro* multi-omics studies have shown that the SARS-CoV-2 dysregulates PI3K/Akt/mTOR and HIF-1 signaling in infected cells. These pathways regulate glycolysis by altering glucose transporters (GLUT) across cell membranes. Targeting these pathways with inhibitors such as MK2206 (Akt inhibitor) or 2- deoxy-D-glucose (2-DG, glycolysis inhibitor) can lower the viral burden in the cells *in vitro* [7, 8]. This opens the area for host-based metabolic strategies to inhibit viruses as an additional way other than direct-acting antivirals to weaken the viral replication by metabolic intervention.

In this study, we performed plasma proteomics targeting 92 plasma proteins related to inflammation and plasma untargeted metabolomics followed by immune phenotyping of the lymphocyte and monocyte cell population towards the metabolite transporters. We also performed quantitative untargeted proteomics studies *in vitro* by infecting lung, liver, kidney, and colon cells with the SARS-CoV-2 virus to understand the viral-induced metabolic rewiring. We also modulate the key metabolic pathways identified in the patient-based metabolomics data and cell-model based quantitative proteomics data to regulate the viral reproduction. Our clinical and experimental studies thus provided an account of metabolic control during SARS-CoV-2 infection that can aid antiviral therapeutics in COVID-19 through metabolic perturbation.

## Results

### Patient characteristics

The study population included healthy controls (HC, n=31), SARS-CoV-2 PCR positive hospitalized-mild (mild, O_2_ consumption<4lit/min, n=29) and hospitalized-severe (severe, O_2_ consumption≥4lit/min, n=12) patients. The mild and severe groups were matched by gender ≥ (male: 79% vs 91%, p=0.6514), BMI [median (IQR): 29 (25-31) vs 28 (25-34); p=0.8622] and age [median (IQR): 57 (44-63) vs 57 (52-69); p=0.2831]. The HC has significantly lower age [median (IQR): 48 (46-55)], lower BMI [median (IQR): 24 (21-25)] (Table S1). The IgG CoV-2 antibody test showed 10 of the HC were CoV-2 antibody positive (HC-CoV-2 Ab+, Fig S1).

Among the COVID-19 patients, the classical co-morbidities were observed at 45% (13/29) in mild and at 66% (8/12) in severe (p=0.3058). The samples were collected within median (IQR) 2 (2-3.5) days of hospitalization [median (IQR) mild: 2 (1-3) and severe 3 (2-4); p=0.1170].

### Plasma proteomics identified distinct clusters of HC and COVID-19 individuals

We performed targeted proteomics analyses (secretome) looking at 92 plasma proteins involved in inflammatory responses. As expected, several cytokines and chemokines were significantly elevated in COVID-19 patients (mild and severe) compared to HCs (HC and HC-CoV-2 Ab+), including IL-6 (Fig 1a). Pathway enrichment analyses of the proteins that were significantly changed between HCs and COVID-19 patients revealed that the majority of altered proteins were involved in cytokine-cytokine receptor interaction and chemokine signaling, followed by intestinal network for IgA production, IL-17 signaling pathway, and Toll-like receptor signaling pathway; to name the top five pathways (Fig 1b). Interestingly, 11 proteins were altered between the mild and severe COVID-19 patients (Fig 1a and 1c): hepatocyte growth factor (HGF), pleiotrophin (PTN), the chemokines CXCL12, CXCL13, and CCL23 (also known as macrophage inflammatory protein 3 or MIP-3), monocyte-chemotactic protein (MCP-3, also known as CCL7), interleukin 12 (IL-12), tumor necrosis factor-like weak inducer of apoptosis (TWEAK), vascular endothelial growth factor A (VEGFA), angiopoietin 2 (ANGPT2), and Fas ligand (FASLG) (adj p<0.05). The majority of these proteins were elevated in COVID-19 patients with highest levels in the severe group, except for IL-12 which was increased in the mild group compared to both severe group and HCs (Fig 1a and 1b). Further, FASLG followed the opposite trend of being lower in COVID-19 patients compared to HC and lowest in severe COVID-19 patients. This data showed that although IL-12 levels were increased in COVID-19 patients compared to HC, the COVID-19 severe patients had reduced IL-12 compared to the mild patients.

**Fig 1.**
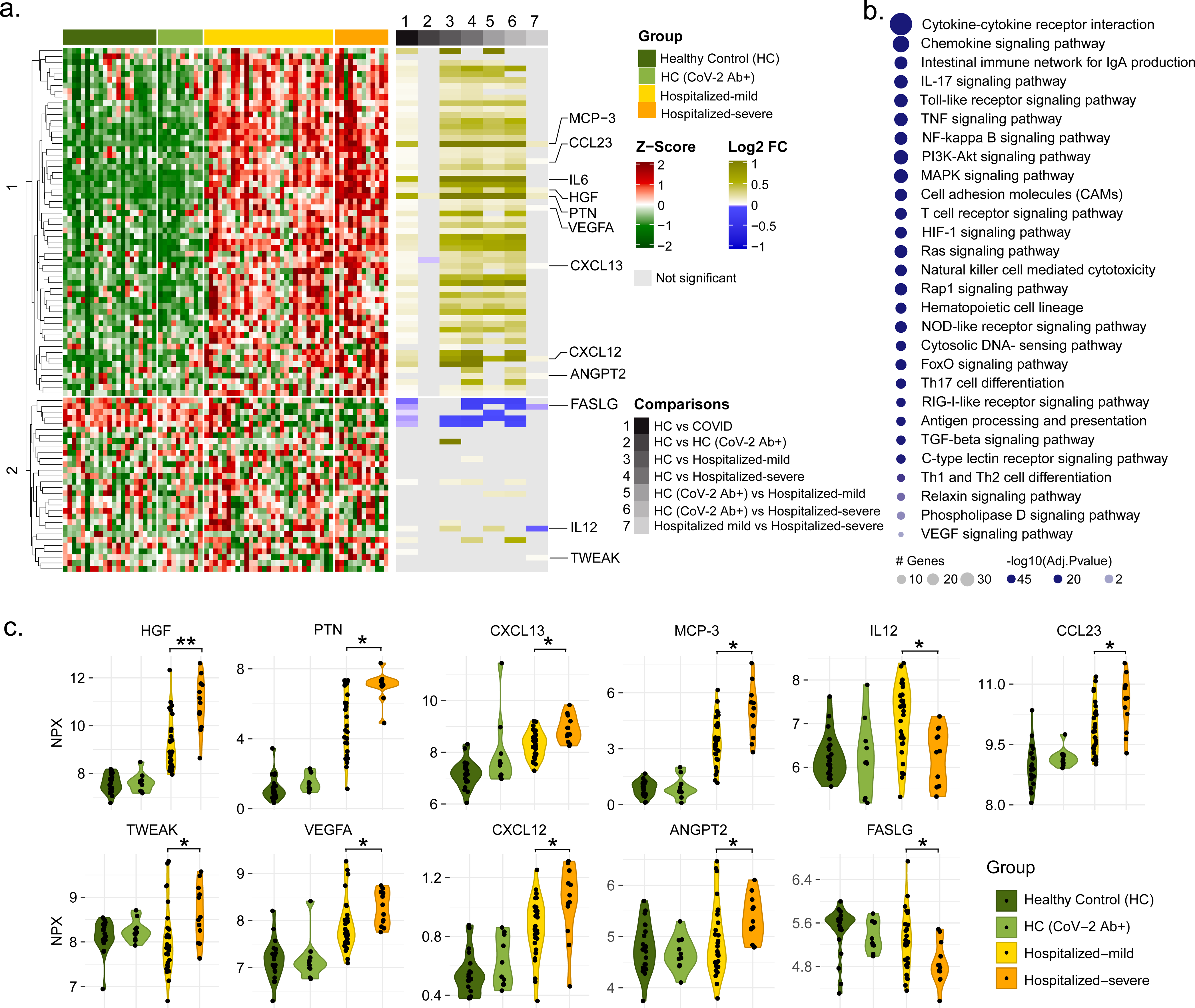
Targeted plasma proteomics in COVID-19 patients. **(a)** Heatmap of Z-score transformed quantitative measurements of all proteins detected by the immuno-oncology panel. Column annotation represents each patient sample and their corresponding groups and pairs of statistical analysis. Rows are proteins hierarchically clustered based on the Euclidean distance and complete linkage method. Names of proteins which are identified as significant in any of the statistical analysis printed. **(b)** KEGG pathway enrichment analysis results of significantly changed proteins between healthy control (HC + HC-CoV-2 Ab+) and COVID-19 (hospitalized mild+hospitalized severe) groups. **(c)** Violin plot of significantly regulated (Mann-Whitney U test) proteins between hospitalized mild and hospitalized severe, *adj p<0.05, **adj p<0.01.

### Distinct amino acid and carbohydrate profile in COVID-19 patients

The plasma metabolomic profile followed a pattern similar to the plasma proteomics. However, no metabolites were significantly different between HC and HC-CoV-2 Ab+ (adj p>0.05). Therefore we combined the two groups as HC for further analysis. The distribution of all samples for metabolite enrichment showed a fair separation between samples of HC and COVID-19 patients (Fig 2a). Differential analysis between COVID-19 patients and healthy individuals after adjusting for age, gender, and BMI identified 444 significantly regulated metabolites (adj p<0.05), many of which are lipids followed by amino acids (Fig 2b). Metabolite set enrichment analysis of the significant metabolites (adj p<0.05) identified amino acid-related pathways were most predominantly affected during infection, as shown in Fig 2c. Hierarchical clustering of the metabolites showed two clusters that had distinct enrichment patterns in COVID-19 infected patients compared to HCs (Fig 2d). Among these, amino acids such as glycine, proline, tryptophan, alanine, histidine, glutamine and arginine had lower levels in COVID-19 patients, while glutamate, aspartate and phenylalanine had higher levels (Fig 2d and S2) as also observed in earlier studies [1, 3, 9]. Interestingly, metabolites of the central carbon metabolism including glycolysis (glucose, 3-phosphoglycerate, pyruvate, lactate) and tricarboxylic acid (TCA) cycle (citrate, aconitate, α-ketoglutarate) showed distinct changes when comparing HCs and COVID-α 19 patients and the different COVID-19 disease states (Fig S3).

**Fig 2.**
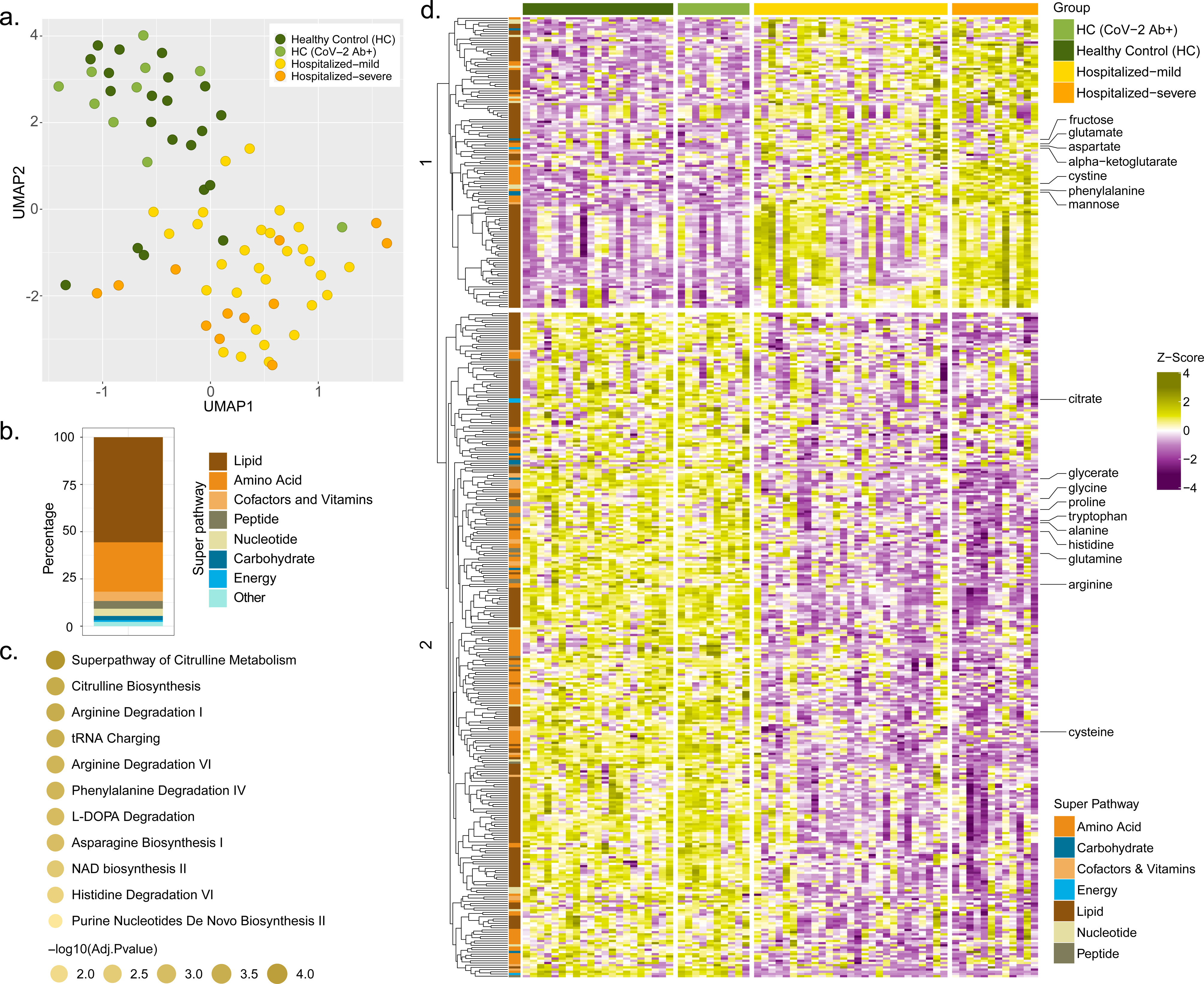
Untargeted global plasma metabolomics in COVID-19 patients. **(a)** Sample distribution for quantitative metabolite measurements plotted in 2-dimensional space after performing dimensionality reduction using UMAP. **(b)** Stacked bar plots visualizing percentage of metabolites significantly changed between healthy control (HC + HC-CoV-2 Ab+) and COVID- 19 (hospitalized mild + hospitalized severe) group concerning their corresponding super-pathways and sub-pathways. **(c)** Metabolic set enrichment analysis using the significantly enriched metabolites between HCs and COVID-19 patients. The size of the bubble indicates adjusted p-values. **(d)** Heatmap of log2 scaled and Z-score transformed measurements of metabolites significantly changed between healthy control (HC + HC-CoV-2 Ab+) and COVID- 19 (hospitalized mild + hospitalized severe) groups. Column annotation represents each patient sample and the corresponding groups. Row annotation represents super pathways of the metabolites. Rows are metabolites hierarchically clustered based on Euclidean distance and complete linkage method.

### Severe COVID-19 patients show a distinct metabolic profile with mannose as a biomarker

Next, we aimed to identify the metabolic signature in COVID-19 severe patients. Statistical analysis found 88 metabolites that significantly differed in the severe group compared to mild samples. Hierarchical clustering of the metabolites showed two clusters with moderately distinct enrichment patterns in severe samples compared to mild samples (Fig 3a). Metabolic pathway enrichment analysis of the significant metabolites by IPA showed that several amino acid-related pathways, IL-12 signaling and production in macrophages and insulin signaling pathways were mainly dysregulated in the severe patient samples compared to mild ones (Fig 3b). Next, we sought to identify biomarkers that differentiate the severe and mild samples. Using R package MUVR that is suitable for small sample size, a total of eight metabolites were identified as biomarkers (Fig S4). After adjusting for age, gender and BMI, seven remained significant of which four had higher abundance and three had lower abundance in COVID-19 severe patients compared to the mild ones (Fig 3c and S4). Mannose was identified as one of the biomarkers of disease severity, being upregulated in COVID-19 infection and also in severe patients compared to mild ones. COVID-19 infection was associated with increased glutamate levels while glucose and mannose were determinants of the severity of the disease (Fig 3d and 3e). This data suggests alterations in the glycolysis/gluconeogenesis, glutaminolysis and mannose metabolism in COVID-19 patients irrespective of severity.

**Fig 3.**
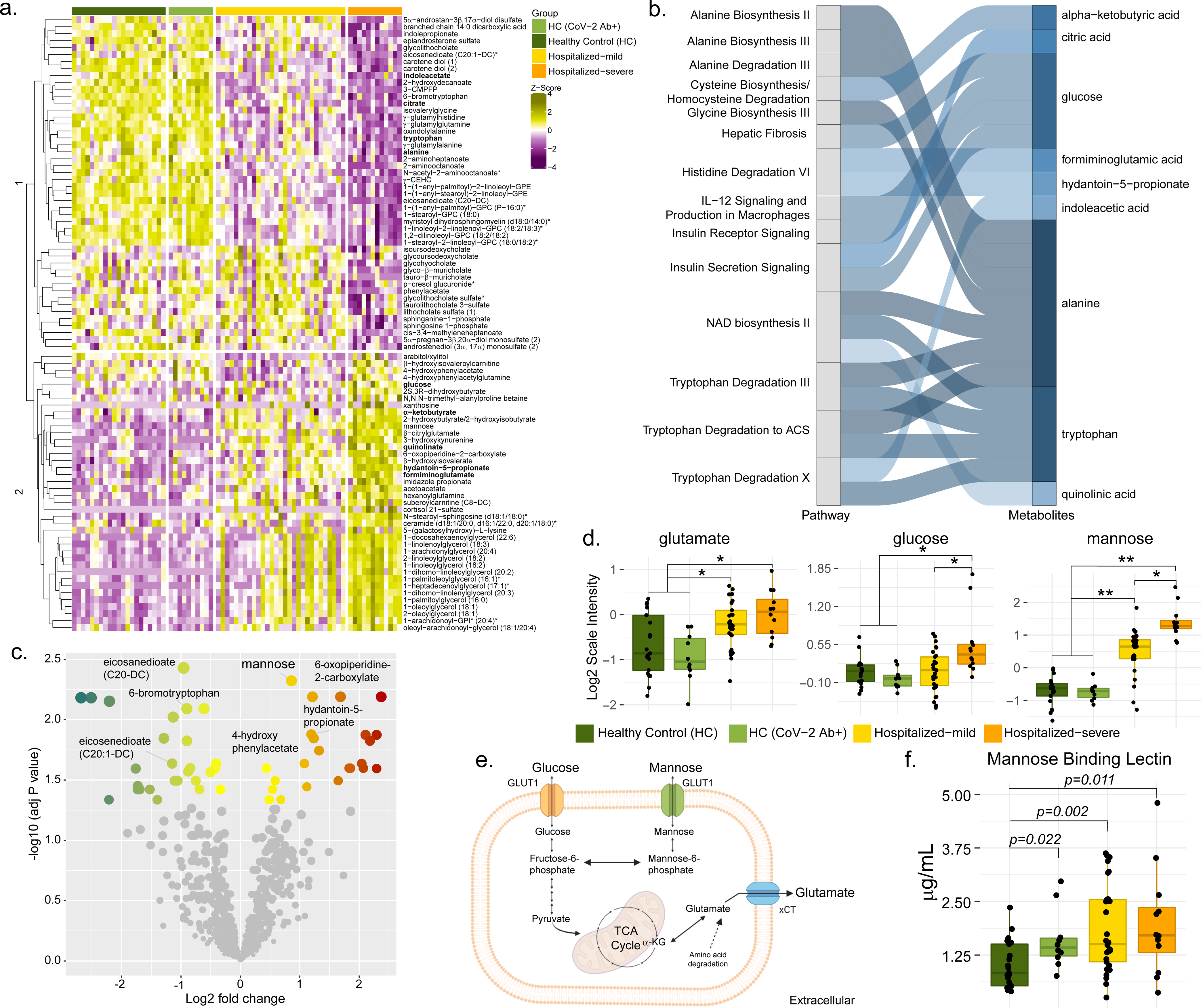
Untargeted plasma metabolites differing between the mild and severe hospitalized COVID19 patients. **(a)** Heatmap of log2 scaled, and Z-score transformed significantly changed metabolites between hospitalized mild and hospitalized severe groups. Column annotation represents each patient sample and the corresponding groups. Rows are metabolites hierarchically clustered based on Euclidean distance and complete linkage method. **(b)** Alluvial plot representing pathways resulted from IPA pathway enrichment analysis using all metabolites that differs significantly between hospitalized mild and severe groups. **(c)** Volcano plot showing all the metabolites that differ significantly between hospitalized mild and hospitalized severe groups. **(d)** Box plots of key metabolites glutamate, glucose and mannose **(e)** Schematic representation of the key steps of glycolysis, mannose and glutamate metabolism and TCA cycle. **(f)** Box plots of soluble mannose-binding lectin levels in patients’ plasma. P-values determined by Mann- Whitney U test.

### Increased mannose-binding lectin (MBL) in COVID-19 patients without any correlation with mannose levels

Plasma mannose can bind to several C-type lectins (*e.g.* MBL) and play an important role in viral pathogenesis by recognizing glycans present in the viral envelope and subsequently activating antiviral immune response and T-cells [10, 11]. Our targeted secretome data identified the C-type lectin receptor signaling pathway as one of the top 28 ranked protein pathways that were significantly changed between HCs and COVID-19 patients (Fig 1b). We, therefore, measured plasma levels of soluble MBL using ELISA and observed an increase in COVID-19 patients compared to HC. Strikingly, there was no significant difference between the mild and severe COVID-19 patients, but the HC-CoV-2 Ab+ individuals showed increased MBL levels compared to HC (Fig 3f). This data shows a prominent elevation of MBL during COVID-19 infection that can persist over a prolonged duration of time after recovery. We did not observe any correlation between MBL and mannose in COVID-19 patients [Spearman correlation: 0.1437 (95%CI: - 0.1806 - 0.4399)]. We, therefore, speculate that elevated MBL might not directly be the consequence of the higher plasma mannose levels but regulated by SARS-CoV-2.

### Immune phenotyping of glucose, glutamate, and mannose transporters

Metabolite transporters are known to dictate immune cell activity by controlling access to nutrients, thereby maintaining cellular homeostasis [12]. Therefore, we next measured the expression of glucose/mannose and glutamate transporters that were associated with SARS-CoV-2 infection or disease severity, GLUT1 (SLC2A1) and xCT (SLC7A11) respectively, in PBMCs of HC (n=19), HC-CoV-2 Ab+ (n=9) and COVID-19 patients: mild (n=21) and severe (n=11) using flow cytometry. The relative frequency of lymphocytes significantly decreased in COVID-19 patients than HCs, which was more prominent in severe patients (Fig 4a). In total lymphocytic populations the CD3^+^ T-cells were significantly reduced in COVID-19 severe patients compared to mild patients and HCs (Fig 4b). Although there was no difference in total monocyte frequencies, we observed a mild increase in the frequency of intermediate monocytes and a significant decrease in non-classical monocytes in the COVID-19 patients than HCs (Fig 4a and 4b). This highlights the potential role of monocytes in COVID19 as was also recently reported in single-cell transcriptomics data [4] and functional analysis on COVID-19 patient monocytes [13].

**Fig 4.**
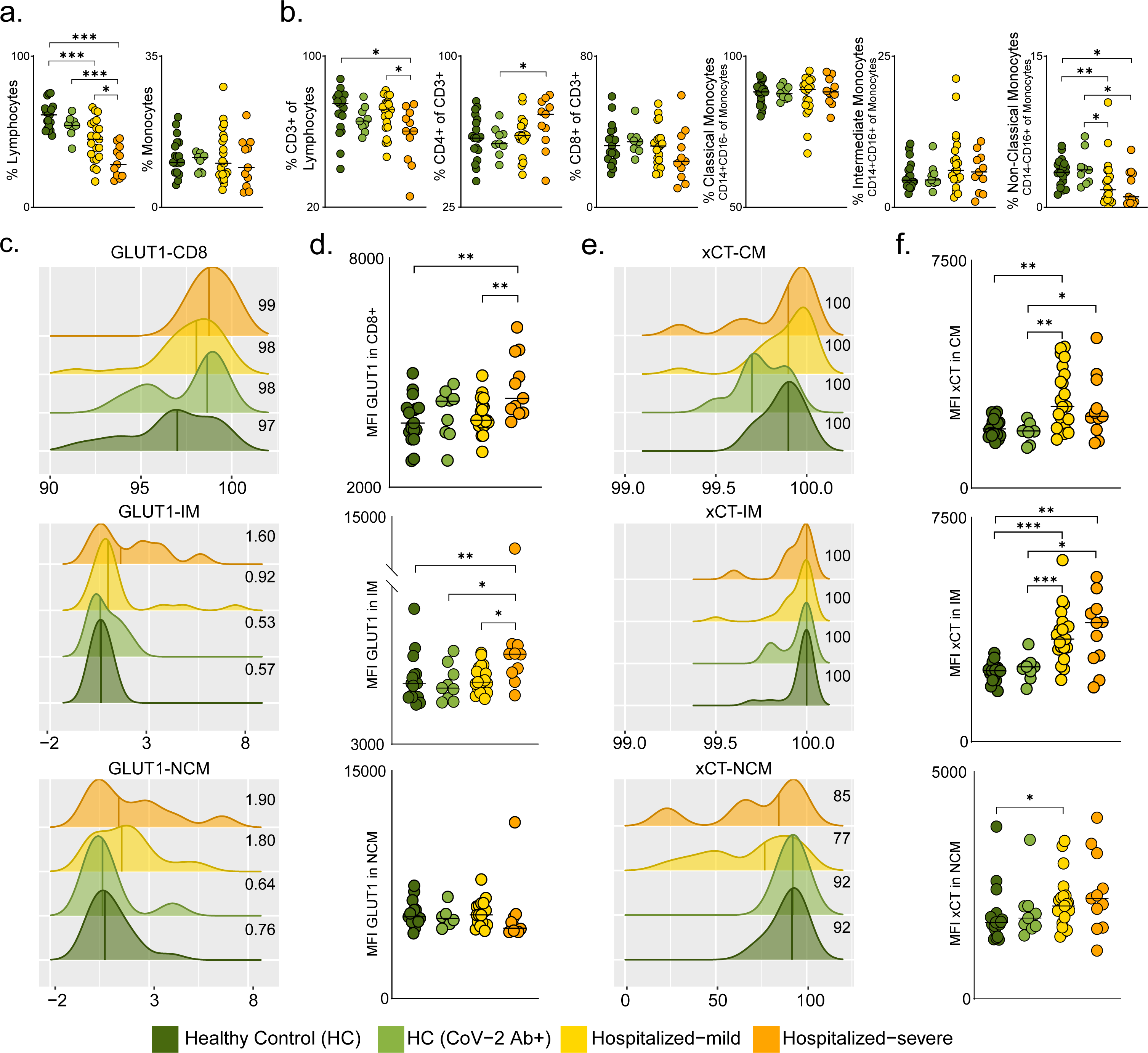
Glucose, mannose and glutamate transporters in COVID-19 severity. **(a)** Percentage of total lymphocytes and monocytes in all 4 patient groups; **(b)** Percentage of PBMC subpopulations, CD3^+^ T-cells of lymphocytes, CD4^+^ T-cells (of CD3^+^ cells) CD8^+^ T-cells (of CD3^+^ cells), classical monocytes (CM, CD14^+^CD16^-^ of monocytes), intermediate monocytes (IM, CD14^+^CD16^+^ of monocytes) and non-classical monocytes (NCM, CD14^-^CD16^+^ of monocytes). Median values are indicated by lines. **(c)** Density plot of percentage of CD8^+^ T-cells, IM and NCM expressing GLUT1. Histograms show percentage of cells expressing GLUT1 (x-axis) and GLUT1 read density of each sample (y-axis). The median percentage of cells expressing GLUT1 is shown for each patient group. **(d)** MFI of GLUT1 in CD8^+^ T-cells, IM and NCM in all four patient groups. **(e)** Density plot of percentage of CM, IM and NCM expressing xCT. Histograms show percentage of cells expressing xCT (x-axis) and xCT read density of each sample (y-axis). The median percentage of cells expressing xCT is shown for each patient group. **(f)** MFI of xCT in CM, IM and NCM in all four patient groups. In all the panels the median values are indicated by lines, p-values are determined by Mann-Whitney U test, *p<0.05, **p<0.01, and ***p<0.001.

More than 98% CD8^+^ T-cells expressed GLUT1, and the surface expression of GLUT1 was significantly higher in COVID-19 severe patients than mild and HCs (Fig 4c and 4d). We also found increased frequencies of intermediate and non-classical monocytes expressing GLUT1 and significantly higher surface expression of GLUT1 on the intermediate monocytes of COVID-19 severe patients compared to HCs (Fig 4c and 4d). While all classical and intermediate monocytes expressed xCT, we observed decreased frequency of non-classical monocytes expressing xCT in COVID-19 patients compared to HCs (Fig 4e). The surface expression of xCT was higher in classical and intermediate subpopulations of COVID-19 patients than HCs (Fig 4f), emphasizing the role of metabolite transporters in monocytes in COVID-19 infection and disease severity.

### SARS-CoV-2 regulates central carbon metabolic pathways in a cell-type-specific manner

Our previous study, together with other observational studies, indicate that SARS-CoV-2 infection causes dysregulation of PI3K/Akt/mTOR and HIF-1 signaling pathway [7, 14, 15], and affect mitochondrial functions [13, 16]. Based on these findings, we hypothesized that the altered extracellular glucose, mannose and glutamate levels are due to dysregulated carbohydrate metabolism and mitochondrial function. Therefore, to identify the acute host cellular response to the SARS-CoV-2 infection we infected different human cell lines including Calu-3 (lung), Caco- 2 (colon), 293FT (kidney) and Huh7 (liver) followed by untargeted quantitative proteomics 24 hours post-infection (hpi) at multiplicity of infection (moi) of 1 [17]. Differential protein abundance analysis identified 6462 proteins differentially expressed in Calu-3, 177 in Caco-2, four in Huh7 and none in 293FT following 24 hpi. This data indicated an acute response predominantly in the lung cells that are the primary site of SARS-CoV-2 infection. The protein set enrichment analysis restricted to metabolic pathways identified that most of the highly abundant proteins in infected Calu-3 cells belonged to pentose phosphate pathway (PPP), fructose and mannose metabolism, as well as amino acid biosynthesis (Fig 5a). Proteins detected at a lower level in the infected cells mainly belonged to TCA cycle, oxidative phosphorylation and N-glycan biosynthesis (Fig 5a). Parallelly, in patients’ metabolomic analysis we observed unbalanced levels of glycolysis, fructose and mannose metabolism and TCA cycle intermediates (Fig 2d and 3d). We, therefore, focused our analysis on the proteins (n=78) that are a part of glycolysis/gluconeogenesis, fructose and mannose metabolism and TCA cycle (KEGG Human 2019) (Fig 5b). A clear change in metabolic poise was observed upon SARS-CoV-2 infection in Calu-3 cells where a majority of the significantly upregulated proteins belonged to glycolysis/gluconeogenesis and fructose and mannose metabolism while most of the proteins of the TCA cycle were significantly downregulated (Fig 5b and S5). However this phenomenon was not observed in the other three cell culture models (Fig S6). Only two out of 177 proteins identified were significantly different in Caco-2 cells (ACSS1 and PFKFB4) and no differences were observed in Huh7 cells out of the four identified proteins in the three pathways mentioned above (Fig S6). This shows that Calu-3 cells, which are lung epithelial cells, have a distinct metabolic modulation caused by SARS-CoV-2 infection. Interestingly, although all the mitochondrial TCA cycle enzymes were downregulated, cytosolic enzymes such as MDH1, IDH1, ACO1 and ACLY that convert TCA cycle intermediates outside the mitochondria were upregulated in infected Calu-3 cells (Fig 5c). This points towards dysfunctional mitochondria caused by COVID-19 infection. Alterations in mitochondrial DNA (mtDNA) copy number in circulating blood cells can serve as a surrogate for mitochondrial dysfunction [18]. Indeed in our patient cohort we observed a decreasing trend of the mtDNA copy numbers with the disease severity (Fig 5d). In addition to changes in glucose and glutamate (Fig 3d), we also observed a significant increase in metabolites such as pyruvate, lactate and α-ketoglutarate (more pronounced in mild patients) and decrease in citrate and aconitate in COVID-19 patients compared to HCs (Fig S6). This indicated an impact of SARS-CoV-2 infection on glycolysis and glutaminolysis to meet biosynthetic and bioenergetic demands. In order to determine the requirement of glycolysis and glutaminolysis for optimal replication of SARS-CoV-2 in Calu-3 cells, we blocked these pathways using 2-DG and 6-diazo-5-oxo-L-norleucine (DON) respectively (Fig 5e) following infection. Infectivity of SARS-CoV-2, quantified as relative *E- gene* levels in cells lysates, showed ∼50-fold decrease in 2-DG treated cells and >100-fold decrease in DON treated cells (Fig 5f). This was also corroborated with virus production in the cell culture supernatant, quantified by viral *E-gene* levels that decreased by more than 2log10 RNA copies/ml in both 2-DG and DON treated cells compared to untreated cells (Fig 5g). While several studies have shown the role of glycolysis on SARS-CoV-2 infection [14, 15], so far there is no direct evidence linking the role of glutaminolysis to replication and spread of SARS-CoV-2 and here we show for the first time that both glutaminolysis and glycolysis are essential for SARS-CoV-2 infection.

**Fig 5.**
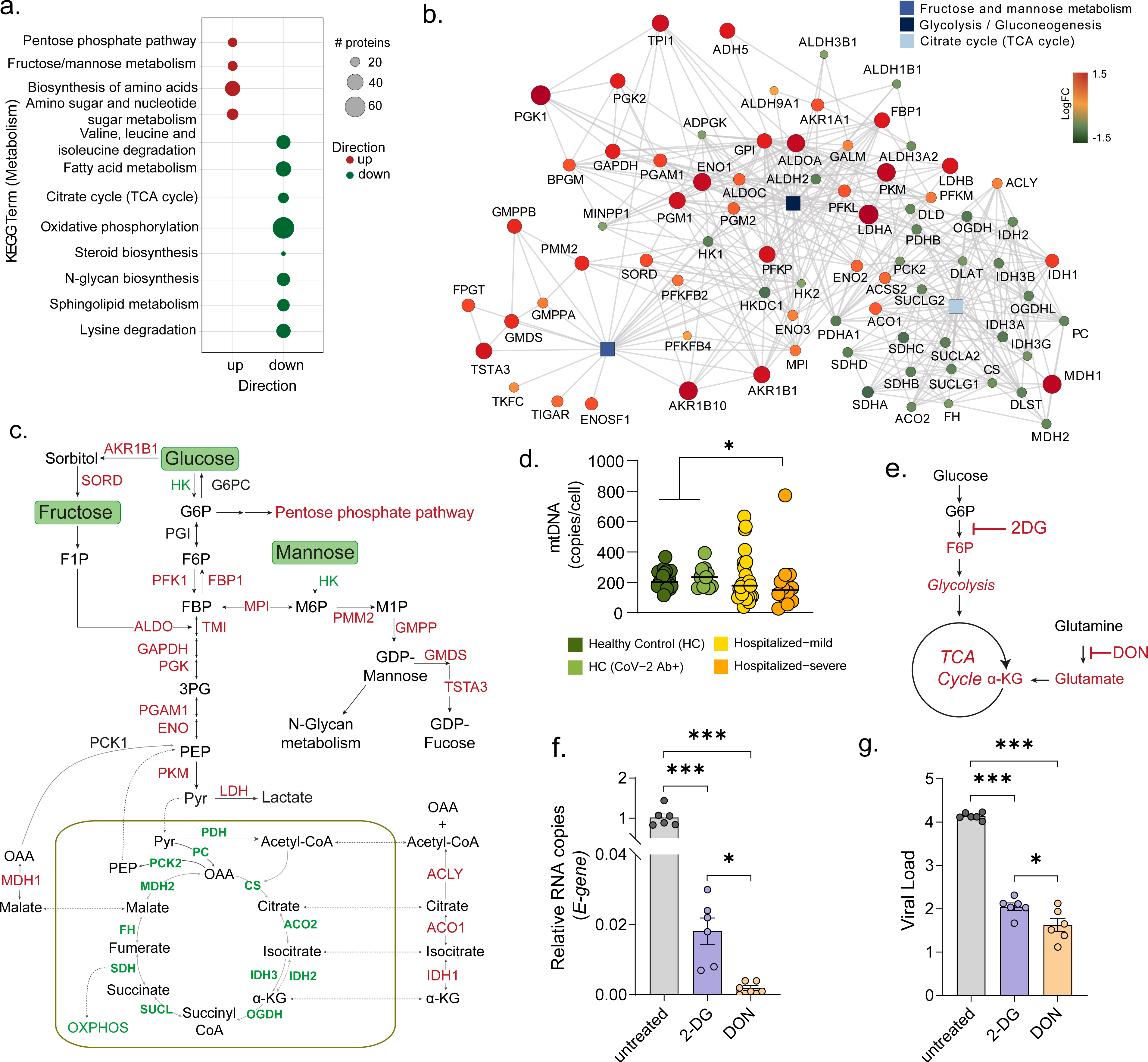
Cell-specific regulation of central carbon metabolic pathways by SARS-CoV-2. **(a)** Bubble plots of protein set enrichment analysis (adj. p<0.05) restricted to metabolic pathways showing highly upregulated (red) and downregulated (green) proteins in SARS-CoV-2 infected Calu-3 cells compared to mock-infected cells. Bubble size is relative to number of proteins. **(b)** Network analysis of proteins from glycolysis/gluconeogenesis, fructose and mannose metabolism and TCA cycle that were significantly different in SARS-CoV-2 infected and mock-infected Calu-3 cells. Rectangular shapes represent the three pathways. Circular shapes show each protein that is either upregulated (red) or downregulated (green) in infected cells compared to mock-infected cells. The size of the circle indicates fold-change. Lines denote connection of each protein to its respective pathway and connection between each protein-protein (STRING, confidence>0.7). **(c)** Schematic map of the glycolysis/gluconeogenesis, fructose and mannose metabolism and TCA cycle. Red indicates significantly upregulated proteins and green indicates significantly downregulated proteins in SARS-CoV-2 infected Calu-3 cells. **(d)** mtDNA copy number in whole blood cells in all 4 patient groups. Median values are indicated by lines, p-values are determined by Mann-Whitney U test, *p<0.05. **(e)** Schematic of inhibitors of metabolic pathways, 2-DG inhibits glycolysis and DON inhibits glutaminolysis. **(f, g)** Viral load of SARS-CoV-2 determined by RT-qPCR targeting the viral *E-gene* is measured in **(f)** cells lysates and **(g)** cell culture supernatants, at moi 0.001 in Calu-3 cells treated with 2-DG or DONas indicated. The data is represented as mean±SEM of two individual experiments, triplicates in each experiment. P-values are determined by student T-test, *p<0.05, ***p<0.001.

### Role of increased sugars in SARS-CoV-2 infection and the complement system in vitro

To understand the role of sugars like glucose and mannose in SARS-CoV-2 infection, we performed *in vitro* infection assays in Calu-3 cells with varying media concentrations of glucose (11.1, 22.2, 44.4mM) and mannose (0, 11.1, 22.2, 44.4mM) with 0.001moi. We did not observe any statistically significant difference in virus production in the supernatant in the glucose/mannose concentrations tested (p>0.05, Fig 6a), while we found a significant reduction in viral *E-gene* expression at cellular level with high glucose concentration of 44.4 mM (Fig 6b, p<0.05). Supplementation with high mannose did not cause any significant change in expression of the *E-gene*. Overall, our data indicate that increased glucose levels but not mannose levels has an effect on viral replication *in vitro*.

**Fig 6.**
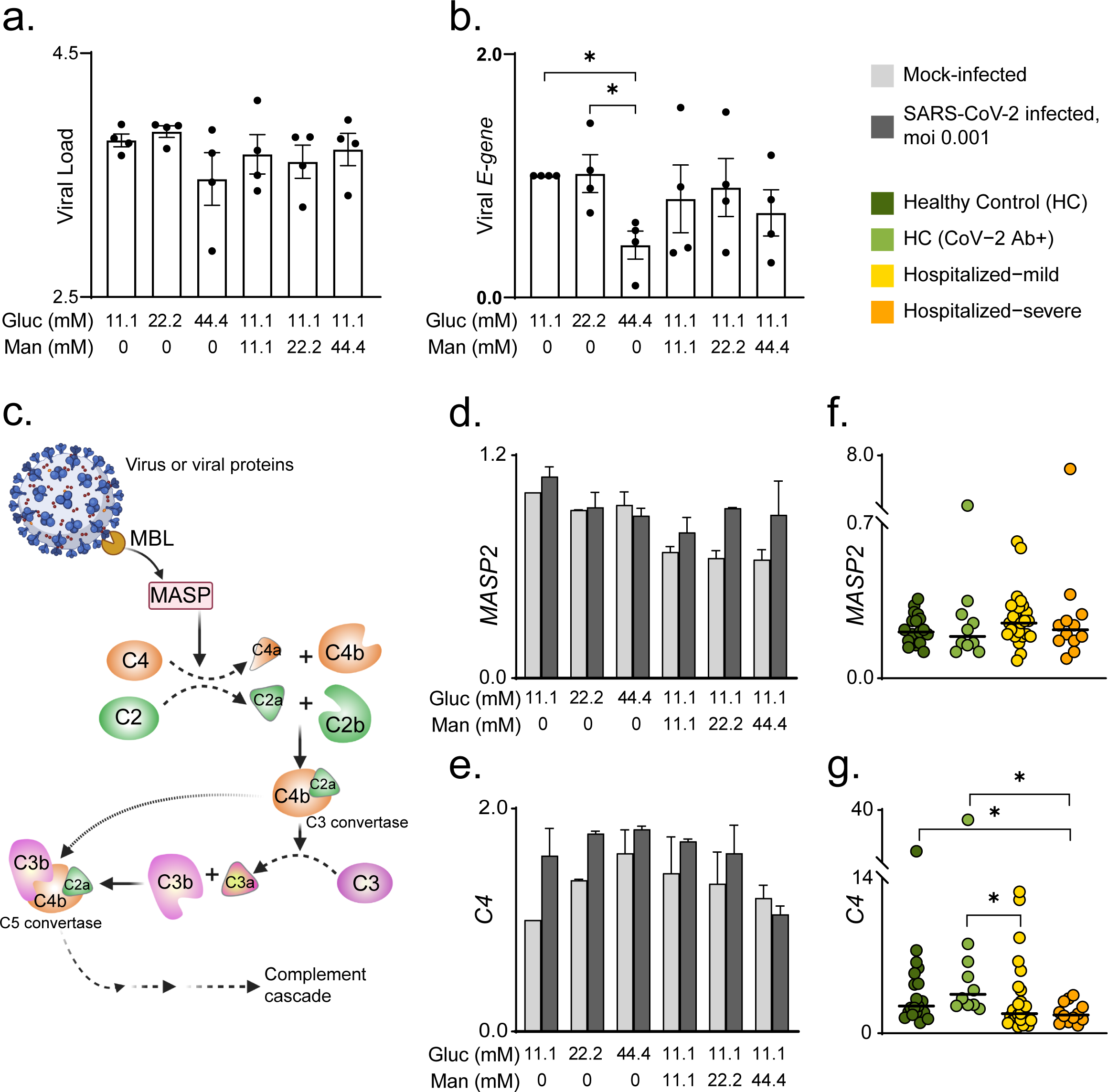
Role of glucose and mannose in SARS-CoV-2 infectivity and complement pathways. Viral load of SARS-CoV-2 determined by RT-qPCR targeting the viral *E-gene* is measured in **(a)** cell culture supernatants, and **(b)** cell lysates at moi 0.001 infection of Calu-3 cells grown in different glucose and mannose concentrations (mM) as indicated. The data is represented as mean±SEM of two independent experiments, duplicates in each experiment. P-values are determined by student T-test, *p<0.05. **(c)** Schematic of complement cascade showing pathway upstream of C3 and C5 activation. Gene expressions of **(d)** *MASP2*, and **(e)** *C4* in SARS-CoV-2 infected (moi 0.001) and mock-infected Calu-3 cells at different glucose and mannose concentrations (mM) as indicated. All gene expression data in Calu-3 cells (viral *E-gene*, *MASP2* and *C4*) at each glucose/mannose concentration were compared with baseline expression at 11.1mM glucose, 0mM mannose and data represented as mean±SEM. Gene expressions of **(f)** *MASP2*, and **(g)** *C4* in whole blood of HC, HC-CoV-2 Ab+, COVID-19 hospitalized mild and hospitalized severe patients determined by RT-qPCR. Median values are indicated by line, p- values are determined by Mann-Whitney-U test, *p<0.05 and **p<0.01.

Several studies, including ours, have shown an effect of SARS-CoV-2 infection on complement and coagulation cascades and thrombosis as unique features of COVID-19 infection [7, 19, 20]. Temporal transcriptomic and proteomic profiling of SARS-CoV-2 infected Huh7 cells have also revealed an upregulation of proteins involved in complement activation at a later stage of infection [20]. Since complement component C4 is part of the cascade that leads to C3 activation in the classical and MBL pathways, we also studied whether increased glucose/mannose in the media during infection affected upstream events of C3 activation (Fig 6c). This was performed by determining gene expression levels of MBL associated serine protease 2 (*MASP2*) and complement *C4* in Calu-3 cells infected with SARS-CoV-2 (moi 0.001) at different concentrations of glucose and mannose. Mannose availability in the culture medium slightly reduced gene expression of *MASP2* in mock-infected that increased upon infection. However no significant changes in *MASP2* expression were observed compared to the basal condition (Fig 6d). An interesting trend was observed with *C4* gene expression (Fig 6e). In mock-infected cells, we observed an increase in *C4* expression with increasing glucose concentrations that decreased dose-dependently following addition of mannose. In the infected cells, a general increase was detected in *C4* expression compared to mock that also followed a similar trend in different glucose and mannose concentrations as was observed in mock-infected cells (Fig 6e). Since COVID-19 patients in this study had high plasma concentrations of glucose and mannose, we also measured *MASP2* and *C4* gene expression in whole blood originating from these patients. No change was observed in *MASP2* expression while a significant decrease in C4 expression was observed in COVID-19 severe patients compared to HCs (Fig 6f and 6g). Combined, *in vitro* and *in vivo* data indicate a potential role of extracellular glucose and mannose concentrations in complement activation. Further, our data suggest that SARS-CoV-2 infection can transcriptionally repress the C4 system in the presence of high mannose, but this needs further mechanistic investigations.

## Discussion

In this study, we used metabolomics, proteomics and immunophenotyping to observe the effect of SARS-CoV-2 infection on metabolic dysregulation in COVID-19 patients and performed *in vitro* infection in four different cell models to find out potential host metabolic regulation during acute SARS-CoV-2 infection. Our study has shed light on the role of monocytes, especially in severe COVID-19 disease. As expected, COVID-19 patients presented a cytokine storm. Interestingly IL-12 plasma levels were decreased in severe compared to mild COVID-19 patients. Among the carbohydrates, plasma mannose emerged as a biomarker for disease severity, but *in vitro* assays showed no effect of mannose on viral replication. Furthermore, host cellular response following SARS-CoV-2 infection identified a strong acute metabolic adaptation in the lung epithelial cells (Calu-3) by modulating central carbon metabolism and indicative of mitochondrial dysfunction that is also observed in severe COVID-19 patients. Glycolysis and glutaminolysis are essential for virus replication and metabolic perturbations of these processes can impede SARS-CoV-2 and could be an attractive antiviral strategy. Finally, SARS-CoV-2 potentially regulates the *C4* system transcriptionally in the presence of carbohydrates.

As reported, the cytokine storm syndrome is evident in COVID-19 patients [21, 22]. Several plasma pro-inflammatory cytokines including IL-6 were elevated in both mild and severe COVID-19 patients. In our study, though we observed higher levels of soluble IL-12 in mild patients, the severe patients showed IL-12 levels similar to HCs. Elevated plasma levels of IL-12 have also been described in hospitalized patients infected with other coronaviruses, such as SARS-CoV and Middle East respiratory syndrome coronavirus (MERS-CoV) [23, 24]. However, to our knowledge, no associations of IL-12 levels with disease severity were reported in these infections. In a recent study on COVID-19 patients on ventilators, the reported low level of IL-12 could be caused by impaired monocytes and affect natural killer (NK) cell functions [25, 26]. *In vitro* studies on IL-12 administration have shown enhanced host cellular responses that generally promote virus clearance and host recovery from infection [27]. IL-12 also plays a critical role in viral immunity by activating the NK cells and promoting differentiation of Th1 CD4^+^ T-cells.

Based on the different levels of IL-12 in COVID-19 patients at varying disease stages, its role in disease severity needs further attention. Both DNA and RNA viruses rewire host cell metabolism by altering central carbon metabolic pathways such as glycolysis, gluconeogenesis, PPP, TCA cycle, amino acid synthesis/degradation, and lipid synthesis. In our metabolomics data, we observed increased glucose, pyruvate and lactate levels in the plasma of COVID-19 patients compared to HCs indicative of toxic metabolic dysregulation during acute phase of infection. Under aerobic, standard growth conditions, primary mammalian cells use glucose for generation of ATP. Glucose is enzymatically broken down to pyruvate that can feed into the TCA cycle in the mitochondria ultimately driving electron transport chain to generate ATP with the help of oxygen.

Viral infections including SARS-CoV-2 are known to enhance the glycolytic flux and increase the production of lactate from pyruvate [6, 28, 29]. Viruses can target glycolysis by regulating glucose transporters’ expression, which is also vital for immune cell activation during host cellular response [30, 31]. Increased GLUT1 does not only result in higher glucose uptake but also gives rise to increased PPP intermediates that enhance nucleotide pool required for viral replication [5]. GLUT1 transports glucose, mannose, glucosamine and docosahexaenoic acid across the cell membrane [32]. We observed a significant increase in surface expression (MFI) of GLUT1 in CD8^+^ T-cells and intermediate monocytes in COVID-19 severe patients. We also measured surface expression of xCT, a cystine/glutamate antiporter that exchanges glutamate for cystine essential for maintenance of redox balance. While there was a decrease in frequency of only non-classical monocytes expressing xCT in COVID-19 patients, a significant increase in surface expression of xCT in classical and intermediate monocytes COVID-19 patients was noted. To the best of our knowledge, the association of xCT expression in respiratory viral diseases has not been studied before and our data for the first time highlights the crucial role of expression of xCT in monocytes. A growing body of evidence highlights the potential role and metabolic status of monocytes in COVID-19 disease severity [4, 13, 33]. Combining all the data, metabolite transporters, xCT and GLUT1, play an essential role in disease severity that could be linked to higher plasma glucose and mannose levels. The specific changes of metabolic transporters were more prominent in monocytes, indicating that metabolic profile of the different monocytic sub-populations could contribute to mediating the severity of the disease.

Plasma mannose emerges as a robust biomarker of disease severity and can also lead to activation of the lectin complement system at a later stage of infection [20]. In addition to mannose, we also observed increased levels of MBL in both COVID-19 patients and convalescent patients compared to the COVID-19 negative individuals. However, no correlation between plasma MBL and mannose were observed in COVID-19 and these two can very well be independent factors. Monomeric mannose is a crucial element of N-linked glycosylation of proteins. Recent studies reported that plasma mannose levels were an indicator of glycogenolysis as well as glucose tolerance and associated with the future risk of developing chronic diseases such as type 2 diabetes, cardiovascular diseases, and albuminuria [34, 35]. In our study cohort, the incidence of type 2 diabetes was low in all study groups and can therefore not explain the high plasma mannose levels in COVID-19 patients. This further strongly suggests that increased mannose is an effect of SARS-CoV-2 infection. However, in light of recent research indicating a possibility of a bidirectional link between SARS-CoV-2 and diabetes, it is tempting to speculate that increased mannose has a role to play in new-onset diabetes after SARS-CoV-2 infection [36–39]. Mannose can bind to several sugar-binding proteins, called lectins, including C-type lectins, such as MBL, mannose receptor, and DC-SIGN. These receptors recognize carbohydrates, particularly on the surface of microorganisms leading to activation of the complements cascade and phagocytosis [10]. Although N-linked mannose residues have been identified on SARS-CoV-2 Spike protein, it is improbable that the elevated plasma mannose levels in the patients would be derived from the virus itself [40]. The processing of endogenous glycoconjugates and their subsequent efflux from the cells are currently thought to be responsible for mannose levels in the blood and steady-state maintenance [41]. Interestingly, a recent study by Heindel et al. describes endogenous high mannose levels as a key mediator of influenza virus-induced pathogenesis and disease severity [42]. High mannose is induced through unfolded protein response pathway and the influenza virus-infected cells are recognized in a high mannose-dependent manner by MBL. Finally, authors state that MBL signaling contributes to disease severity through complement cascade activation and inflammatory response. High mannose and/or high MBL could thus dysregulate the immune system and lead to severe damage associated with disease severity [42]. Activation of the complement is one of the features seen in COVID-19, as described in earlier studies [7, 43]. In concordance with that and with Heindel *et al*., we also noticed increased plasma MBL in COVID-19 and even in healthy convalescent controls compared to HC. This increase in MBL could potentially regulate the complement cascade.

In the *in vitro* Calu-3 infection model, most of the proteins from carbohydrate metabolism and PPP were upregulated while most of the proteins of TCA cycle, oxidative phosphorylation, and fatty acid metabolism were downregulated in infected cells compared to the mock-infected controls (Fig 5). Further delineation of the pathways indicated an inefficient mitochondrial metabolism as majority of the TCA cycle enzymes were downregulated in the infected cells. This was in line with the decreased mtDNA copy numbers in severe COVID-19 patients (Fig 5f) indicating a possible mitochondrial dysfunction as reported previously [13, 44–46].

The metabolism and concentration of sugars and amino acids such as glucose, mannose, glutamine and glutamate among others, play an important role in cellular metabolic homeostasis and are targeted by viruses for their replication. Recent studies have shown that elevated glycolysis favors SARS-CoV-2 infection and replication [14, 15]. While glutaminolysis has been implicated as a carbon source for other human DNA and RNA viruses [5]. Our data shows for the first time that glutaminolysis is also crucial for SARS-CoV-2 infection and replication. The inhibition of glutaminolysis has a larger effect on viral replication and production compared to the inhibition of the glycolysis in lung cell model (Fig 5f and 5g). Glutaminolysis is a process of converting glutamine to TCA cycle intermediates and also essential for biosynthesis of proteins, lipids and nucleic acids. Some viruses (e.g. Herpes simplex virus 1, human cytomegalovirus, Hepatitis C virus etc) use glutamine as an anaplerotic substrate to replenish TCA cycle via generation of α-ketoglutarate [29, 47]. Recently, researchers have proposed that the metabolic α reprogramming of glutamine in SARS-CoV-2 can trigger pathogenesis. They further hypothesized that metabolic intervention of glutaminolysis could be an antiviral strategy for COVID-19 [47–49]. Although the exact underlying mechanism is unknown, our *in vitro* study shows that SARS-CoV-2 replication depends on both glycolysis and glutaminolysis.

Finally, to elucidate the effects of extracellular glucose and mannose in both infection of SARS-CoV-2 and its impact on the complement cascade, we established an *in vitro* infection set up with varying media concentrations of glucose and mannose post-infection. Virus production in the cell culture supernatant was unaffected by both glucose and mannose concentrations. However, high extracellular glucose decreased viral infectivity, measured as relative expression of viral *E-gene* in cells. This contradicts the earlier finding that an increase in glucose concentration aids in virus replication [14]. However, it is to be noted, the observations by Codo et al. were made in peripheral monocytes while we performed our experiments in Calu-3 cells. Despite this fact, several studies attempted to isolate infectious virus particles from blood but failed [50, 51] and we were not able to find any residual viral RNA in patients’ whole blood cells. Also, blood cell populations do not express the ACE2 or TMPRSS2 (Blood Atlas in the Human Protein Atlas [52]). Thus the infectivity of the blood cell population including lymphocytes and monocytes needs careful consideration [53].

In conclusion, our patient based multi-omics studies and *in vitro* analysis emphasizes the need to understand the host metabolic reprogramming due to acute SARS-CoV-2 infection. Among other factors, the role of carbohydrate and amino acid transporters, mainly in the monocytic- macrophage lineages, under the altered central carbon metabolism regulated by AKT/mTOR/HIF-1 signaling may potentially define disease severity. The metabolic alteration in glucose, mannose, lactate, pyruvate, and glutamate in severe COVID-19 cases need further clinical considerations. Changes in these metabolites might have a sustained effect on insulin resistance, type 2 diabetes, neurocognitive impairments, and multiorgan failure which is already reported in COVID-19 infection.

## Methods

### Experimental Model and Subject Details

#### Study designing, patients

The COVID-19 patients (n=41) who were PCR positive and hospitalized in May 2020, were recruited from the South Hospital, Stockholm. Based on the oxygen requirements the patients were categorized into, 1) Hospitalized-mild (O_2_ consumption <4lit/min) and 2) Hospitalized-severe (O_2_ consumption ≥4lit/min). The exclusion criteria ≥ included known liver cirrhosis, severe renal insufficiency (estimated eGFR≤30mL/min/1.73m^2^), ≤ chronic obstructive pulmonary disease, and chronic lung disease leading to habitual SpO_2_ 92%. ≤ Additionally, COVID-19 PCR negative samples (HC herein, n=31) were also collected. IgG antibody was tested on the HC samples as described previously [51] and ten samples turned out to be SARS-CoV-2 Ab positive further defined as HC-CoV-2 Ab+. The study was approved by regional ethics committees of Stockholm (dnr 2020-01865). All participants gave informed consent. The patient identity was anonymized and delinked before analysis.

#### Cell lines and viruses

Human colon adenocarcinoma cell line, Caco-2, and lung adenocarcinoma cell line, Calu-3 (ATCC^®^ HTB-55^™^), were purchased from CLS Cell Lines Service GmbH, Germany and LGC Standards, UK, respectively. Hepatocyte-derived carcinoma cell line, Huh7, was obtained from Marburg Virology Lab, Philipps-Universität Marburg, Marburg, Germany matching the STR reference profile of Huh7 [54], and human embryonic kidney cell line, 293FT (Invitrogen). SARS-CoV-2 virus used in this study was the first virus isolated from a Swedish patient[55].

### Method Details

#### Materials

All information regarding reagents, antibodies, and critical commercial kits are listed in Table S2.

#### IgG Antibody detection against SARS-CoV-2

In brief, 96-well ELISA plates (Nunc MaxiSorp, ThermoFisher Scientific) were coated with SARS-CoV-2 N protein, diluted 1:1000 in 50 mM sodium carbonate pH 9.6 (1 ug/ml final concentration), at 4°C overnight. The plates were then blocked for 1h using PBS containing 1% BSA and 2% goat serum (dilution buffer) at 37°C. Serum samples were serially diluted from 1/200 to 1/6400 in dilution buffer and incubated on the plates at 37°C for 1h. Antibodies to the N protein was detected by incubation for 1h at 37°C with anti-human IgG peroxidase (1:30,000, Sigma). The plates were washed three times with PBS with 0,05% Tween-20 between each step. The bound conjugate was visualized using Tetramethylbenzidine (Sigma, USA) substrate. The OD was measured at 450 nm with subtraction of background at 650 nm using a TECAN Infinite M200 plate reader (Tecan, USA).

#### Plasma inflammation profiling and metabolomics

The plasma inflammation profiling was performed using proximity extension assay technology targeting 96 inflammation markers by Olink Immuno-Oncology panel (Olink, Sweden). Plasma untargeted metabolomics was performed by Global Metabolomics (HD4) in Metabolon, NC, US as described by us recently [56]. The metabolomics method is ISO 9001:2015 certified and the lab is accredited by the College of American Pathologist (CAP), USA.

#### Statistical and bioinformatics analysis

For targeted proteomics data analysis, we used Mann-Whitney U through the R package stats v3.6.1 for pair-wise analysis as the data was not normally distributed. For metabolomics data, dimensionality reduction of all samples were performed with Uniform Manifold Approximation and Projection (UMAP) using R package umap v0.2.6.0 [57]. Reduced dimensions of the data were plotted in 2D space using R package ggplot v3.3.2[58]. The metabolite measurements were log2 scaled before differential analysis. Differential analysis was done using R/Bioconductor package limma v3.42.2 [59]. R package MUVR v0.0.973 [60] was used for biomarker discovery. It is a software package that performs predictive multivariate modeling by integrating minimally biased variable selection procedure into repeated double cross-validation architecture. Random forest core modeling was selected from the package for biomarker identification. Minimal-optimal variables selected by the model were considered as biomarkers. Correlation analysis was performed using corr.test function from the package psych v1.9.12.31 based on Spearman rank correlations. Untargeted protein raw data abundance was first filtered for empty rows and quantile normalized. Differential expression analysis was performed with R package limma v3.42.2 2 [59] to determine proteins with differential abundance.

Functional analysis of the proteins was performed using enrichr module of python package GSEAPY v 0.9.16 (https://pypi.org/project/gseapy/) [61, 62], where all the quantified proteins were considered as background. KEGG 2019 Human gene-set library downloaded from Enrichr web resources was used for the enrichment test for molecular pathway analysis. Functional analysis of the metabolites was carried out using Ingenuity Pathway Analysis (IPA) software package. All reported p-values were corrected (*Benjamini*-*Hochberg) throughout* and considered statistically significant if <0.05 unless otherwise stated.

#### Data visualization

Heatmaps were generated using R/Bioconductor package ComplexHeatmap v2.2.0 [63]. Violin plots, box plots, bubble plots and volcano plots were made using geom_violin, geom_boxplots, geom_point objects from R package ggplot2 v3.3.2 respectively. A density plot was created using the function stat_density_ridges function from the R package ggridges v0.5.2. Alluvial plot was made with geom_alluvium function from ggalluvial v0.11.3 R package. Correlation pairs plot was made using ggpairs function from GGally v2.0.0 R package. Network input files were made using R 3.6.3. The network was represented using Cytoscape ver 3.6.1 (https://cytoscape.org/). Protein-protein interactions were retrieved from STRING Db (v5.0) (https://string-db.org/). Only interactions with high confidence (interaction score>0.7) from databases and experiences were kept.

#### Flow Cytometry

Peripheral blood mononuclear cells (PBMCs) were subjected to flow cytometry analysis. Samples were thawed in 37°C water bath and washed with flow cytometry buffer (PBS+2% FBS+2mM EDTA). All samples were stained with Live/Dead fixable near IR dye (Invitrogen), and cell surface markers were detected by incubating cells with relevant antibodies for 20min on ice in flow cytometry buffer (antibodies listed in Table S2). All cells were fixed with 2% paraformaldehyde before acquiring a BD FACS Symphony flow cytometer (BD Bioscience). Compensation setup was performed using single-stained controls prepared with antibody-capture beads: Anti-Mouse Ig, κ/Negative Control Compensation Particles Set (BD κ Biosciences) for mouse antibodies, AbC™ Total Antibody Compensation Bead Kit (Invitrogen) for rabbit antibodies and ArC™ Amine Reactive Compensation Bead Kit (Invitrogen) for use with LIVE/DEAD™ Fixable dead cell stain kits. Flow cytometry data were analyzed and compensated with FlowJo 10.6.2 (TreeStar Inc), Prism 8 (GraphPad Software Inc).

#### Measurement of mtDNA copy number

Mitochondrial DNA (mtDNA) copy number was measured using an Absolute Human Telomere Length and Mitochondrial DNA Copy Number Dual Quantification qPCR Assay Kit (ScienCell Research Laboratories, USA). Each 15µl qPCR reaction contained 7.5µl QuantiNova SYBR green (Qiagen, Sweden), 1µl single copy reference (SCR) and mitochondrial primers, 0.1µl ROX (passive reference dye), 1.9µl DNA/RNA free water and 5µl (1ng/µl) template DNA. Thermal cycle profile comprised incubation at 50°C for 2min and 95°C for 10min before running 40 thermal cycles (95°C for 15s, 54°C for 45 s and 72°C for 45s). mtDNA and SCR primers were run on separate plates and each plate contained a serially diluted DNA sample to calculate PCR efficiency. A reference genomic DNA was added on each plate with known mtDNA copy number (925 copies). Each sample was run in duplicates, and relative mtDNA copy number and SCR to reference was calculated by ΔCT (CT target sample - CT reference sample), after adjusting PCR efficiency using the Pfaffl method [64].

Finally mtDNA copy number per diploid cell of target sample to reference sample was calculated by (2^- ΔΔCT x 925), where ΔΔCT is ΔCT mtDNA/ ΔCT SCR.

#### SARS-CoV-2 infection and proteomics

In 6-well plate, Calu-3 cells were grown for 72h, and Huh7, 293FT, and Caco-2 cells were grown for 24h in DMEM-high glucose (Sigma-Aldrich, USA) supplemented with 10% FBS (Gibco, USA). The cells were either mock-infected in medium only or infected with SARS-CoV-2 at a multiplicity of infection (moi) of 1 in DMEM- high glucose supplemented with 5% FBS. After 1h the inoculum was removed and was replenished with 2mL of fresh DMEM-high glucose containing 5% FBS. Twenty-four hours post-infection (hpi) the supernatant was removed, and the cells were scrapped in 1mL PBS, centrifuged and the washed cell pellets were lysed in 100µL of 2% SDS-lysis buffer (50mM Tris-Cl, 150mM NaCl, 2% SDS and 1mM EDTA) freshly supplemented with protease inhibitor cocktail, phosphatase inhibitor and 1mM DTT. The lysates were heated at 92°C for 10min to deactivate the virus, followed by sonication in a water-bath sonicator for 2min to clear the lysate. Lysates were centrifuged at 13,000rpm for 15min and supernatants were collected. Protein estimation was performed by Bio-rad DC protein assay kit (Bio-Rad Laboratories, USA). The supernatant proteins were used for in-solution digestion and TMT-pro labeled proteomics as described by us previously [7].

#### Metabolic perturbation and virus infection

Calu-3 cells were seeded in 24-well plate, and after 72h of seeding, the cells were infected with SARS-CoV-2 at moi of 0.001 for 1h. Following infection, the cells were treated with DMEM (Gibco, USA) which contained pyruvate (1mM) and glutamine (4mM) as the basal carbon source and were supplemented with 5% FBS and different concentrations of glucose (11.1mM, 22.2mM and 44.4mM) (Gibco, USA) and keeping basal glucose concentration at 11.1mM, different concentrations of mannose (11.1mM, 22.2mM and 44.4mM) (Sigma-Aldrich, USA). To inhibit glycolysis and glutaminolysis, following 1hpi (moi 0.001) the cells were treated with 2-deoxy-D-glucose (2-DG, 10mM) and diazo-5-oxo-L-norleucine (DON, 200µM) respectively. The supernatants were collected after 24hpi and the cells were lysed in TRI reagent (Zymo Research, USA) and stored in -70°C for RNA extraction.

### RT-qPCR Analysis

The virus production and infectivity were determined by qRT-PCR targeting the viral *E-gene* in the supernatant and RNA extracted from the cells. RNA was extracted using Direct-zol™ RNA Miniprep kit according to manufacturer’s instructions (Zymo Research, USA). The supernatant or RNA were directly used for one-step RT-qPCR using PrimeDirect™ Probe RT-qPCR Mix (TaKaRa, Japan) according to manufacturer’s instructions. Primer and probe sequences for the viral *E-gene* and human *RNaseP* gene are listed in Table S3. To measure gene expression of *MASP2* and *C4* in patient blood, whole blood was collected in Tempus™ Blood RNA Tubes (Applied Biosystems, USA) and RNA was extracted using Tempus™ Spin RNA Isolation Kit (Invitrogen, USA). Quality and concentration of extracted RNA was measured using Nanodrop ND-2000 (Thermo Scientific, USA).

The RNA purified from cells and whole blood was reverse transcribed using a High-Capacity cDNA reverse transcription kit (Applied Biosystems, USA) according to manufacturer’s instructions. qPCR reactions were performed using KAPA SYBR Fast qPCR kit (KAPA Biosystems, USA) on an Applied Biosystems™ 7500 Fast qPCR machine. Detailed information on primers is included in Table S3.

### Plasma MBL Measurement

MBL levels in patient plasma were determined using Human MBL Quantikine ELISA Kit (R&D Systems) according to manufacturer’s instructions. The optical density of each well was determined using NanoQuant Infinite M200 plate reader (Tecan, USA) microplate reader at 450nm with background subtraction at 570nm.

### Data and Code Availability

The scaled normalised metabolomics data can be obtained from the dx.doi.org/ 10.6084/m9.figshare.13336862 Proteomics data can be obtained from the ProteomeXchange Consortium via the PRIDE partner repository with the dataset identifier PXD022847.

All the codes are available at github: https://github.com/neogilab/COVIDOMICS

## Acknowledgments

The authors would like to thank Elisabet Storgärd and Ronnie Ask, Study Nurses, *Södersjukhuset* for their excellent support with patient recruitment and all the clinicians and nurses who are the frontline warriors fighting against COVID-19. Authors acknowledge support from the Proteomics Biomedicum; Karolinska Institute, Solna, for LC-MS/MS analysis.

## Supplementary Files

Table S1. Clinical Features of the study population

Table S2: List of reagents, kits and antibodies

Table S3: List of primer and probe sequences

Fig S1: The IgG Ab showed 10 of the HC were CoV-2 Ab-positive.

Fig S2: Metabolite profile of amino acids altered in COVID-19 patients. Line within box plots represents median values, *p<0.05, **p<0.01, ***p<0.001.

Fig S3: Levels of metabolites related to glycolysis/gluconeogenesis and fructose and mannose metabolism and the TCA cycle

Fig S4: Biomarker of the COVID-19 severity identified by MVUR. The size of the bubble indicates rank. Box plot of the biomarkers indicating the level in HCs and COVID-19 patients.: update

Fig S5: Differential protein abundance in Calu-3 cells following SARS-CoV-2 infection after 24h. The analysis was restricted to glycolysis/gluconeogenesis, fructose and mannose metabolism and the TCA cycle.

Fig S6: Differential protein abundance in Caco-2, Huh7 and 293FT cells following SARS-CoV- 2 infection after 24h. The analysis was restricted to glycolysis/gluconeogenesis, fructose and mannose metabolism and the TCA cycle.

Fig S7: Gating strategy of flow cytometry data.

