## Supplementary for "Implications of central carbon metabolism in SARS-CoV-2 replication and disease severity"

### Supplementary Files

#### Supplementary Tables

**Table S1.** Clinical features of study population

| Parameter | HC | Hospitalised-Mild | Hospitalised Severe | P values* | P values** |
|---|---|---|---|---|---|
| number | 31 | 29 | 12 | | |
| Age, years; median (IQR) | 48 (46-55) | 57 (44-63) | 57 (52-69) | 0.0277 | 0.2831 |
| Gender, Male, n (%) | 24 (77%) | 23 (79%) | 11 (91%) | 0.557 | 0.6514 |
| BMI, Median (IQR) | 24 (21-25) | 29 (25-31) | 28 (25-34) | 0.0016 | 0.8622 |
| Sample collection post hospitalisation, days, median (IQR) | - | 2 (1-3) | 3 (2-4) | | 0.1170 |
| Comorbidities, yes (%) | - | 13 (45%) | 8 (66%) | | 0.3058 |
| Obesity, yes (%) | | 2 (7%) | 1 (8%) | | 1 |
| Type 2 Diabetes, yes (%) | | 2 (7%) | 1 (8%) | | 1 |
| Hypertension, yes (%) | | 7 (24%) | 4 (33%) | | 0.7011 |
| Asthma, yes (%) | | 4 (14%) | 3 (25%) | | 0.3978 |

*within the groups, **between hospitalized mild and severe

**Table S2**: List of reagents, kits and antibodies

| | | |
|---|---|---|
| **Antibodies** | | |
| CD4-BUV395-SK3 | BD Biosciences | 563552 |
| CD8-APC-RPA-T8 | Biolegend | 301014 |
| CD14-BV510-M5E2 | Biolegend | 301842 |
| CD3-BV711-OKT3 | Biolegend | 317328 |
| CD16-BV786-3G8 (RUO) | BD Biosciences | 563690 |
| GLUT1-FITC-# 202915 | R&D Systems | FAB1418F |
| xCT-AF594 | Novus Biologicals | NB300-318AF594 |
| CD8-PerCP-HIT8a | Biolegend | 300922 |
| CD3-FITC-OKT3 | Biolegend | 317306 |
| Anti-Human IgG Peroxidase | Sigma-Aldrich | A0170 |
| SARS-CoV-2 N protein | Varnaitė et al., 2020 | GenScript |
| Tetramethylbenzidine | Sigma-Aldrich | T0440 |
| **Critical Commercial Assays** | | |
| Olink Immuno-Oncology | Olink Technology, Sweden | Immuno-Oncology Panel |
| Global Metabolomics (HD4) | Metabolon. Inc., US | HD4 |
| Human MBL Quantikine ELISA Kit | R&D systems | DMBL00 |
| Anti-Mouse Ig, κ/Negative Control Compensation Particles Set | BD Biosciences | 552843 |
| AbC™ Total Antibody Compensation Bead Kit | Invitrogen | A10513 |
| ArC™ Amine Reactive Compensation Bead Kit | Invitrogen | A10346 |
| DMEM-high glucose | Sigma-Aldrich, USA | D6429-500ml |
| Bio-rad DC protein assay kit | Bio-Rad Laboratories, USA | #5000116 |
| TaqMan Fast Virus 1-Step Master Mix | ThermoFisher Scientific | 4444434 |
| Direct-zol™ RNA Miniprep Kit | Zymo Research | R2051 |
| Tempus™ Blood RNA Tubes | Applied Biosystems | 4342792 |
| Tempus™ Spin RNA Isolation Kit | Invitrogen | 4380204 |
| PrimeDirect™ Probe RT-qPCR Mix | TaKaRa, Japan | RR600B |
| KAPA SYBR Fast qPCR kit | Roche | KK4602 |
| TMTpro 16plex Label Reagent set | ThermoFisher Scientific | A44520 |
| Absolute Human Telomere Length and Mitochondrial DNA Copy Number Dual Quantification qPCR Assay Kit | ScienCell Research Laboratories | #8958 |
| QuantiNova SYBR® Green PCR Kit | Qiagen | 208054 |

**Table S3**: List of primer and probe sequences

| **Oligonucleotides** | | |
|---|---|---|
| E_Sarbeco_F1-5'-ACAGGTACGTTAATAGTTAATAGCGT-3' | WHO | |
| E_Sarbeco_R2-5'-ATATTGCAGCAGTACGCACACA-3' | WHO | |
| E_Sarbeco_Probe-5'-[FAM]-ACACTAGCCATCCTTACTGCGCTTCG-[BBQ650]-3' | WHO | |
| RNAseP-F-5'-AGATTTGGACCTGCGAGCG-3' | CDC | |
| RNAseP-R-5'-GAGCGGCTGTCTCCACAAGT-3' | CDC | |
| RNAseP-Probe-5'-[FAM]-TTCTGACCTGAAGGCTCTGCGCG-[BHQ1]-3' | CDC | |
| C4-F-5'-GGGGCCCCACGTCCTGCTGTAT-3' | IDT | (Sorensen et al., 2009) |
| C4-R-5'-CTGCGCTCGGGGTTGTAGTAGTCG-3' | IDT | (Sorensen et al., 2009) |
| MASP2-F-5'-TATGAAAAGCCACCCTATCCA-3' | IDT | (Sorensen et al., 2009) |
| MASP2-R-5'-TGCCCCTCCGCTGTCAC-3' | IDT | (Sorensen et al., 2009) |
| Actin-F-5'-GAGGGAAATCGTGCGTGACA-3' | IDT | (Khan et al., 2016) |
| Actin-R-5'-AATAGTGATGACCTGGCCGT-3' | IDT | (Khan et al., 2016) |

Supplementary Figures

| Pt ID | Plate no. | Mean (1:200) | Mean (1:400) | Mean (1:800) | Mean (1:1600) | Antibody status |
|---|---|---|---|---|---|---|
| HC-03 | 1 | 2,97 | 2,43 | 1,470 | 0,840 | POS |
| HC-10 | 1 | 3,00 | 2,70 | 1,768 | 0,995 | POS |
| HC-12 | 2 | 3,00 | 2,32 | 1,370 | 0,756 | POS |
| HC-14 | 2 | 2,70 | 1,65 | 0,937 | 0,526 | POS |
| HC-15 | 2 | 3,00 | 3,00 | 3,000 | 3,000 | POS |
| HC-18 | 2 | 2,69 | 1,52 | 0,859 | 0,427 | POS |
| HC-23 | 3 | 3,00 | 2,82 | 1,825 | 1,120 | POS |
| HC-26 | 3 | 2,87 | 2,00 | 1,119 | 0,581 | POS |
| HC-13 | 2 | 0,62 | 0,36 | 0,178 | 0,098 | POS |
| HC-32 | 3 | 0,19 | 0,10 | 0,063 | 0,043 | POS |
| HC-21 | 3 | 0,16 | 0,09 | 0,051 | 0,042 | NEG |
| HC-06 | 1 | 0,16 | 0,08 | 0,045 | 0,029 | NEG |
| HC-17 | 2 | 0,15 | 0,09 | 0,053 | 0,038 | NEG |
| HC-31 | 3 | 0,14 | 0,07 | 0,046 | 0,037 | NEG |
| HC-22 | 3 | 0,13 | 0,07 | 0,043 | 0,036 | NEG |
| HC-24 | 3 | 0,12 | 0,07 | 0,045 | 0,034 | NEG |
| HC-25 | 3 | 0,14 | 0,06 | 0,042 | 0,030 | NEG |
| HC-01 | 1 | 0,10 | 0,05 | 0,037 | 0,024 | NEG |
| HC-02 | 1 | 0,06 | 0,03 | 0,023 | 0,019 | NEG |
| HC-04 | 1 | 0,09 | 0,05 | 0,033 | 0,025 | NEG |
| HC-05 | 1 | 0,03 | 0,02 | 0,015 | 0,020 | NEG |
| HC-07 | 1 | 0,05 | 0,03 | 0,019 | 0,014 | NEG |
| HC-08 | 1 | 0,08 | 0,04 | 0,028 | 0,021 | NEG |
| HC-09 | 1 | 0,07 | 0,04 | 0,026 | 0,020 | NEG |
| HC-11 | 2 | 0,08 | 0,05 | 0,032 | 0,124 | NEG |
| HC-16 | 2 | 0,08 | 0,05 | 0,032 | 0,023 | NEG |
| HC-19 | 2 | 0,05 | 0,03 | 0,026 | 0,019 | NEG |
| HC-20 | 2 | 0,06 | 0,04 | 0,028 | 0,023 | NEG |
| HC-27 | 3 | 0,10 | 0,07 | 0,042 | 0,033 | NEG |
| HC-28 | 3 | 0,07 | 0,04 | 0,032 | 0,027 | NEG |
| HC-29 | 3 | 0,07 | 0,04 | 0,029 | 0,024 | NEG |
| Considered Ab+ ≥ | | 0,17 | 0,11 | 0,08 | 0,07 | |

**Fig S1**: The IgG Ab showed 10 of the HC were were CoV-2 Ab-positive.

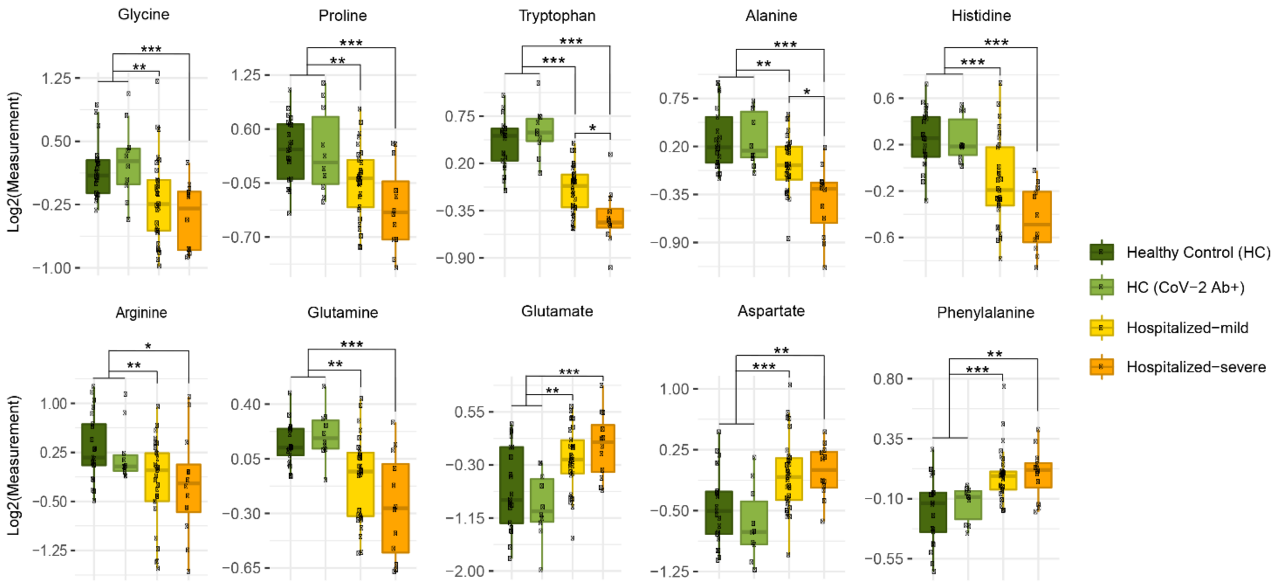


**Fig S2:** Metabolite profile of amino acids altered in COVID-19 patients. Line within box plots represents median values, *p<0.05, **p<0.01, ***p<0.001.

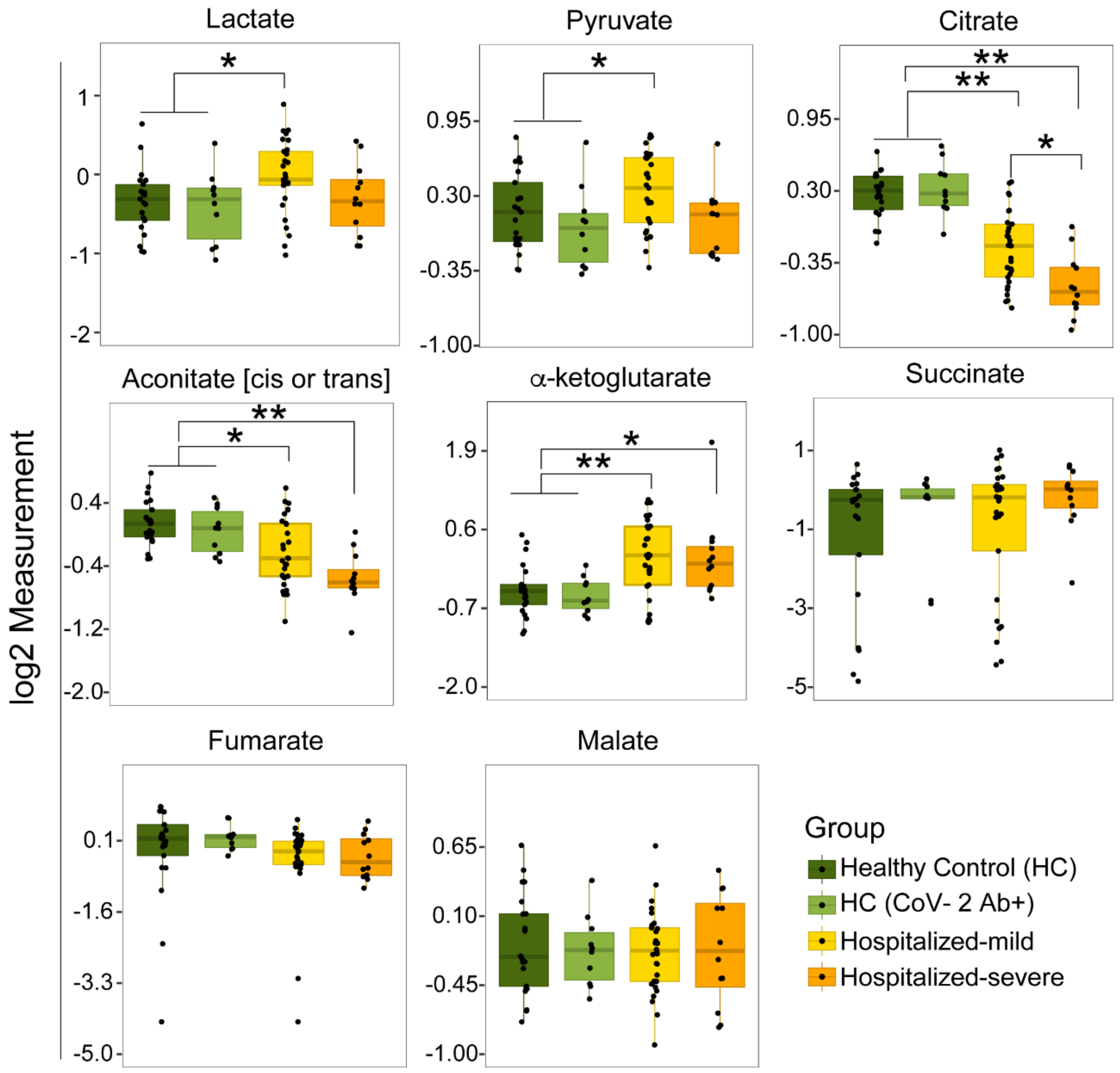


**Fig S3:** Levels of metabolites related to glycolysis/gluconeogenesis and fructose and mannose metabolism and the TCA cycle

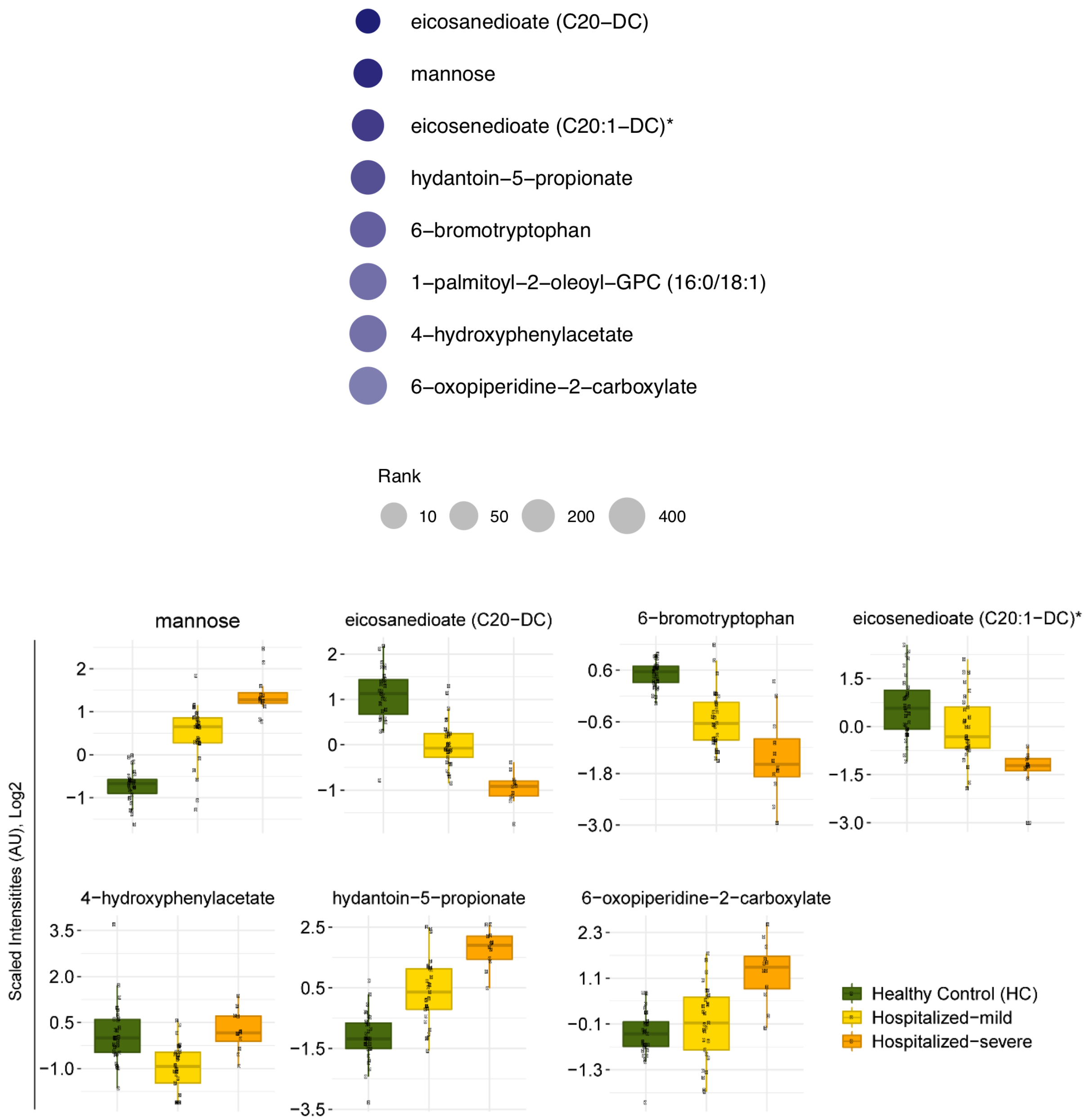


**Fig S4:** Biomarker of the COVID-19 severity identified by MUVR. Size of the bubble indicates rank. Box plot of the biomarks indicating the level in HCs and COVID-19 patients.

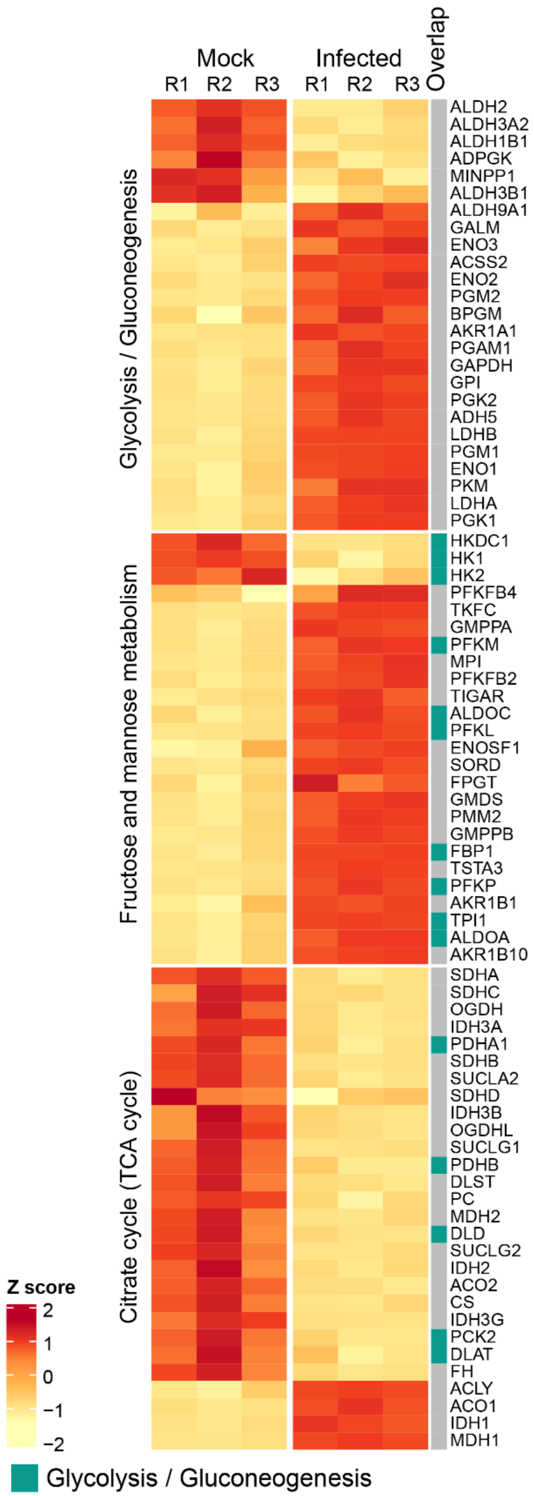


**Fig S5:** Differential protein abundance in Calu-3 cells following SARS-CoV-2 infection after 24 hrs. The analysis was restricted to glycolysis/gluconeogenesis, fructose and mannose metabolism and TCA cycle

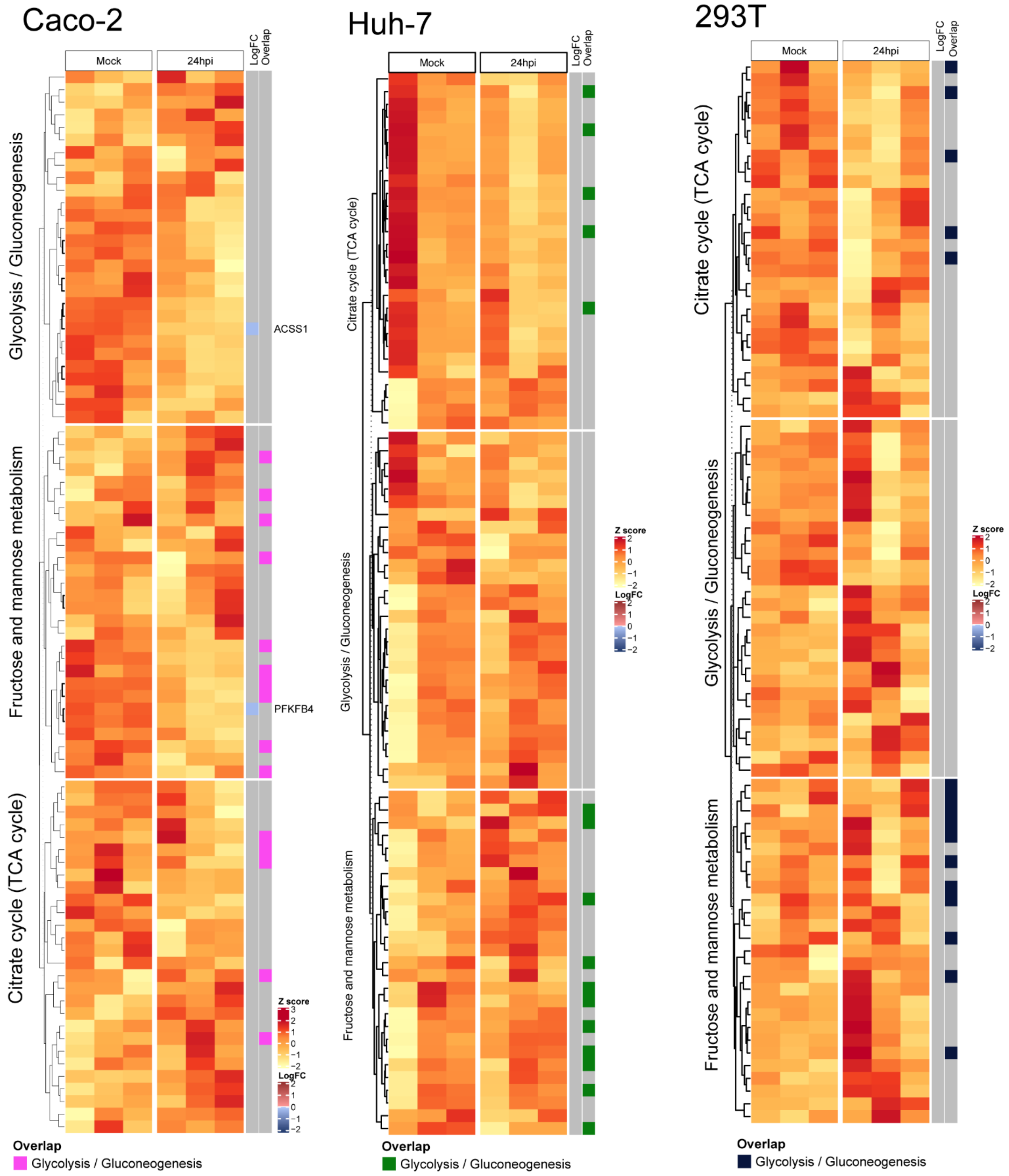


**Fig S6:** Differential protein abundance in Caco-2, Huh7 and 293T cells following SARS-CoV-2 infection after 24 hrs. The analysis was restricted to glycolysis/gluconeogenesis, fructose and mannose metabolism and TCA cycle.

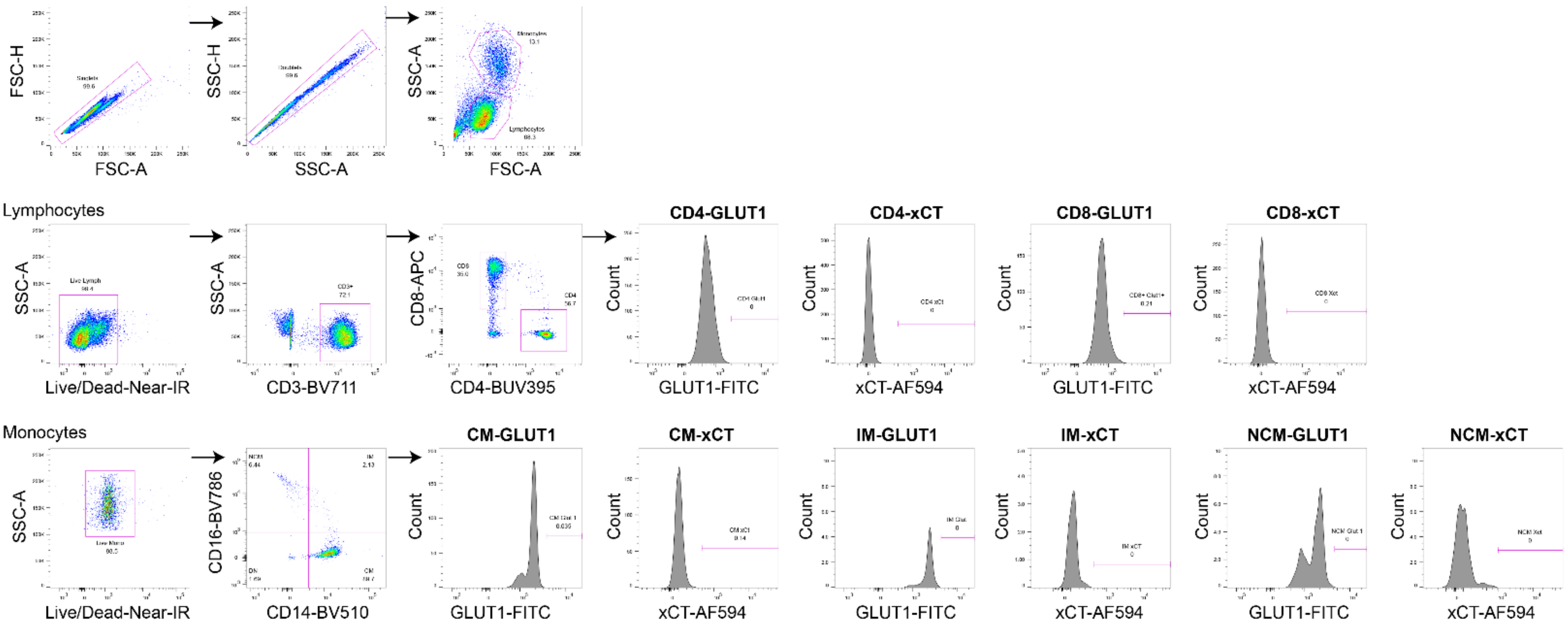


**Fig S7:** Gating strategy of flow cytometry data
